# Hippocampal delta oscillations entrain neuronal activity, modulate gamma amplitude and convey information about running speed on a treadmill

**DOI:** 10.1101/2022.01.24.477542

**Authors:** Alan MB Furtunato, Rafael Pedrosa, Bruno Lobão-Soares, Adriano BL Tort, Hindiael Belchior

## Abstract

Locomotion has long been associated with rhythmic oscillations in the rat hippocampus. Running speed and acceleration affect the spectral density at the theta (6-10 Hz) and gamma (30-150 Hz) bands and the rhythmic entrainment of neuronal activity. However, less is known about other oscillatory rhythms. Recent studies have shown that oscillatory activity in the delta (1-4 Hz) band also relates to locomotion in stationary conditions, such as running on a treadmill or in a running wheel. To further investigate the effects of stationary running on hippocampal oscillations, we recorded CA1 local field potentials and neuronal activity while rats ran at different speed protocols on a treadmill. We found a remarkable oscillatory activity at 2 Hz that was strongly modulated by running speed. Delta power and peak frequency were highest at the fastest running speed, both in constant and progressively increasing speed protocols. Delta and theta oscillations co-occurred and showed independent relationships in their instantaneous power and frequency. Moreover, the delta phase modulated the amplitude of low-gamma (20-50 Hz) oscillations and the spiking activity of putative pyramidal neurons in a speed-dependent manner. Finally, spectral components in the delta frequency range, but not theta, predicted running speed using a naive Bayes classifier. In summary, our study shows that locomotion-related delta oscillations convey information about stationary running speed and coordinate hippocampal rhythmic activity in a speed-dependent manner.

**Statement of significance:** Theta and gamma oscillations are hallmarks of rhythmic activity in the rat hippocampus during locomotion and exploratory behaviors, but less is known about other oscillatory rhythms. This study shows the emergence of delta (∼2 Hz) oscillations in the dorsal CA1 of rats running on a treadmill. Locomotion-related delta oscillations are independent of the concomitant theta activity and convey information about running speed. Delta waves modulate low-gamma oscillations and the spiking activity of pyramidal neurons in a speed-dependent way. These results suggest that oscillatory activity in the delta band aids rhythmic coordination during stationary locomotor behaviors.

## INTRODUCTION

Locomotion has been extensively associated with prominent rhythmic activity in the hippocampal local field potentials (LFP) of freely moving animals since pioneering studies in the 1960s (Vanderwolf and Heron, 1964; Vanderwolf, 1969, 1971; Whishaw and Vanderwolf, 1973; McFarland et al., 1975). Theta (6-10 Hz) oscillations emerge throughout arousal epochs and voluntary movements to coordinate the activity of higher-frequency oscillations and neuronal assemblies in the hippocampus and other interconnected areas (Green and Arduini, 1954; O’Keefe and Recce, 1993; Bragin et al., 1995; Buzsáki, 2002, p. 2005; Siapas et al., 2005; Andersen et al., 2007; Scheffer-Teixeira et al., 2012; Lopes-Dos-Santos et al., 2018). Theta power and frequency positively correlate with running speed in open-field, linear track, and other behavioral tasks (Sławińska and Kasicki, 1998; Shin and Talnov, 2001; Montgomery et al., 2009; Hinman et al., 2011; Richard et al., 2013; Wells et al., 2013; Belchior et al., 2014). Meanwhile, studies examining stationary running on treadmills or running wheels and passive movements in ambulatory apparatus (such as car and train models) reached similar conclusions but also reported significant reductions in theta-running speed relationships and the temporal organization of neuronal activity within theta cycles (Shin and Talnov, 2001; Maurer et al., 2005; Terrazas et al., 2005; Cei et al., 2014; Li et al., 2014; Drieu et al., 2018).

In contrast, delta (1-4 Hz) oscillations are widely expressed in neocortical and subcortical regions and often involved in the synchronization of large neuronal populations typically observed during slow-wave sleep and early stages of development (Siapas and Wilson, 1998; Steriade et al., 2001; Hirase et al., 2001; Sirota et al., 2003; Khazipov and Luhmann, 2006; Emmons et al., 2017). In the rat hippocampus, delta oscillations dominate the LFP during immobility and quiet behaviors - such as licking, chewing, face-washing, scratching, and staying still - but are ceased by sensory stimuli or at the transition from immobility to locomotion, becoming replaced by the theta rhythm (Buzsáki, 2002, 2015; Andersen et al., 2007).

Many reports have corroborated and further highlighted the dichotomy between quiet and active behaviors respectively exhibiting delta and theta oscillations in the hippocampus (Csicsvari et al., 1998, 1999; Czurkó et al., 1999; O’Neill et al., 2006; Montgomery et al., 2009; Allen et al., 2011; Caixeta et al., 2013; Patel et al., 2013; Schultheiss et al., 2020). Nevertheless, provocative new evidence points to the occurrence of hippocampal delta oscillations during stationary locomotion (Furtunato et al., 2020; Safaryan and Mehta, 2021), which suggests that the traditional dichotomy between theta and delta states may not be clear-cut. Corroborating this possibility, here we report an increase in hippocampal delta power and frequency associated with increasing running speed on a treadmill. Locomotion-related delta oscillations modulate higher-frequency oscillations and neuronal activity and also convey information about running speed. In concert with the theta rhythm, the emergence of delta oscillations may provide an alternative mechanism to organize hippocampal activity during running in the presence of static external landmarks.

## MATERIAL AND METHODS

### Subjects

We used six male adult Wistar rats (Rattus *norvegicus*; 3-6 months old; 250-350 g), which were maintained on a 12-hour dark/light cycle (lights on at 07:00 h), individually housed on propylene cages (41 × 34 × 16 cm) with food and water *ad libitum*. Experimental procedures were approved by the Animal Research Ethics Committee of UFRN (permit number 061.069/2017) and were in accordance with the Brazilian legislation for the use of animals in research (Law number 11,794/2008).

### Apparatus and training

We used an electrical treadmill (AVS Projetos) with belt dimensions of 40 × 13.5 cm (Figure 1A). A metallic grid positioned at the back end of the belt was used to deliver low-intensity electrical shocks (0.1 to 0.7 mA at 60 Hz until 1 second) when animals moved out of the belt during the training phase. Electrical shocks were not used during the recording phase. The treadmill speed was automatically controlled by an Arduino Uno board.

**Figure 1.**
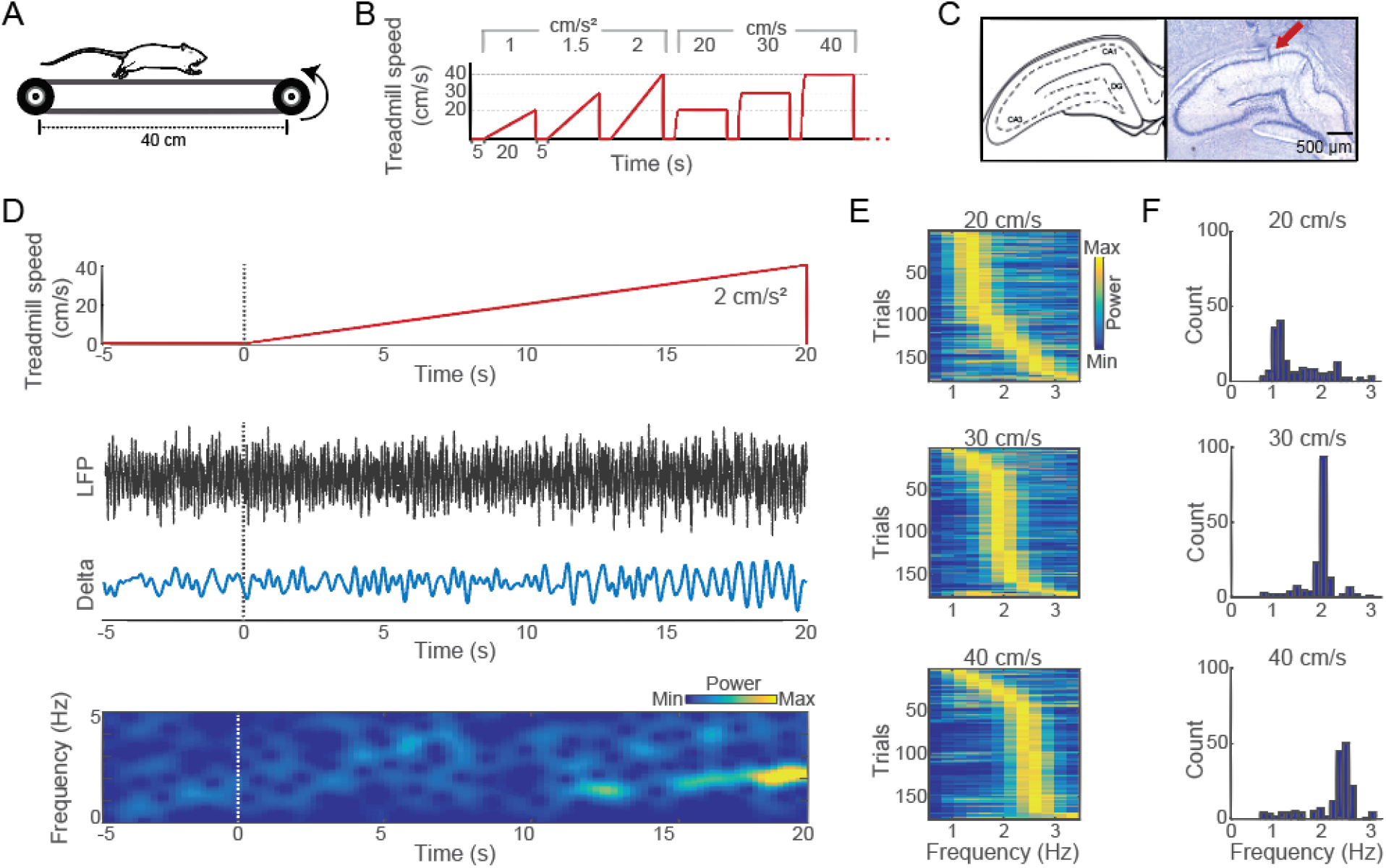
Hippocampal delta oscillations emerge during stationary running. (A) Schematic representation of the treadmill. (B) The treadmill running protocol was composed of 8 blocks of 3 runs at accelerated (1, 1.5, and 2 cm/s²) speeds and 3 runs at constant (20, 30, and 40 cm/s) speeds for 20 seconds interleaved by 5 seconds of intertrial intervals (one block is depicted). (C) Schematic illustration (left) (Paxinos and Watson, 2007) and a representative Nissl-stained coronal section of the dorsal hippocampus (right). The red arrow indicates an electrolytic lesion at the tetrode tip. (D) Treadmill speed (top), raw and delta-filtered (0.5-3.5 Hz) LFPs (middle), and spectral decomposition (bottom) during a 5-s intertrial interval and the following run of an accelerated speed protocol (2 cm/s²). (E) Normalized power spectral density in the delta band for each trial at constant speeds (20, 30, and 40 cm/s), as labeled. (F) Distribution of delta peak frequency values for all 176 running trials at each constant speed protocol.

Animals were trained to execute 48 20-s runs at three acceleration rates (1, 1.5, and 2 cm/s²) and three constant speeds (20, 30, and 40 cm/s). Before the first training session, animals were individually habituated to the treadmill for 3 days. On the first habituation day, rats were placed on the treadmill for 30 minutes with the electrical shock function deactivated and the treadmill turned off. On the second day, the electrical shock function was activated and the treadmill was still off. The same procedure was repeated on the third day, but after 30 minutes of habituation, the treadmill was turned on at low speed (<10 cm/s) for 5 minutes with the electrical shock activated. During the training sessions, the treadmill speeds were set on a maximum of 10, 14, and 18 cm/s and were gradually increased at 2.5 cm/s/day steps until the animals reached the target speeds of 20, 30, and 40 cm/s. The treadmill speed, the duration of the task, and the electrical shock intensity were similar to previous studies (Albeck et al., 2006; Liu et al., 2008; O’Callaghan et al., 2009).

### Surgery

Experimental subjects were submitted to a surgical procedure to chronically implant 3D-printed microdrives of movable tetrodes in the dorsal hippocampus. The rats were anesthetized with isoflurane (5% for induction 5% and 1.5% for maintenance) for 5 minutes, and then received ketamine (100 mg/kg, i.p.) and xylazine (8 mg/kg, i.p.). At the start and the end of the surgical procedure, lidocaine chlorhydrate (3 mg/kg, s.c.) was administered on the cranial surface. The animals were positioned in a stereotaxic apparatus (Kopf Instruments) and subjected to a craniotomy according to the coordinates of the dorsal hippocampus (4.16 mm posterior and 2.4 mm lateral to bregma; Paxinos and Watson, 2007). Six stainless steel screws were implanted in the cranial bones to provide mechanical support for the microdrive, and two additional screws were implanted in the occipital bones to serve as an electrical ground. After the craniotomy, the microdrive was positioned and fixed with acrylic resin. At the end of the surgery, the animals received ∼1 ml glycated saline (5%, i.p.), and a 3-day treatment with enrofloxacin (10 mg/kg, i.p.) and meloxicam (2 mg/kg, s.c.). The animals’ health and overall behavior were supervised during a week of recovery.

### Tetrode positioning

Microdrives supporting 8 and 16 tetrodes were used (Souza et al., 2018). Two animals were implanted with 16-tetrode bilateral microdrives (rats 1 and 3) and the other four animals were implanted with 8-tetrode unilateral microdrives (rats 2, 4, 5, and 6). Tetrodes of nickel-chromium or tungsten microelectrodes (12.7-μm diameter, Sandvik Heating Technology or California Fine Wires Company) were electroplated in gold solution with carbon nanotubes to reduce impedance to ∼100 kOhms at 1 kHz using NanoZ (Neuralynx, according to Tenent et al., 2009). Tetrodes were lowered 1.5 mm into the brain tissue at the end of the surgery and daily steps of 0.2-0.5 mm after surgery recovery until reaching the CA1 area of the dorsal hippocampus. The rats were maintained in a sleep box during the adjustments of tetrode positioning. The proximity to the pyramidal layer was based on electrophysiological signatures of the LFPs, such as the presence of sharp-wave ripples during slow-wave sleep episodes (Buzsáki, 2002, 2015). After each recording session, tetrodes were progressively adjusted in ∼50 μm for the detection of putative new neurons in the next session. Tetrode adjustments were performed >12 h before recording sessions.

### Recording sessions

After surgery recovery, animals were re-trained on the treadmill running until they reached the performance of >90% of complete runs (i.e., without touching the metallic grid). On all the recording sessions, rats were submitted to an exercise protocol of 48 running trials of 20 seconds interspersed by 5 seconds of rest (Figure 1B). The exercise protocol consisted of eight trials of six runs, in which the first 3 runs were performed at the acceleration rates of 1, 1.5, and 2 cm/s² followed by 3 runs at the constant speeds of 20, 30, and 40 cm/s. Rats were maintained on the treadmill and recorded for 5 minutes before the running protocol started.

Electrophysiological signals were obtained using one or two 32-channel headstages (RHD2132, Intan Technologies) with an Open Ephys acquisition system. The acquisition system and the treadmill were electrically grounded. The raw electrophysiological signals were digitized, pre-amplified (20x), and sampled at 30 kHz. Digital video recordings were made at 30 frames/second using a webcam (C920, Logitech) perpendicularly positioned to the treadmill. The treadmill was controlled by the same computer used for video and electrophysiological recordings; data were synchronized through TTL events and stored for posterior analyses. The experimental subjects contributed unequally to the total of 22 recording sessions; rats 1, 2, 3, 4, 5, and 6 respectively contributed with 2, 1, 5, 6, 6, and 2 sessions.

### Euthanasia and craniotomy

At the end of the experiments, the animals were anesthetized in a pressurized box with isoflurane (induction 5%, maintenance 1.5%) for approximately 5 minutes followed by ketamine (100 mg/kg, i.p.) and xylazine (8 mg/kg, i.p.). Then, an electrical current (0.05 µA for 30 s) was applied to each electrode to cause micro lesions at the brain tissue around the tips of the electrodes (Insight Equipamentos). The animals were intracardially perfused with 200 ml of saline phosphate buffer solution (37°C and pH 7.4) and 300 ml of buffered paraformaldehyde (37°C and pH 7.4). The brains were extracted and maintained in buffered paraformaldehyde solution for 76 hours, and then for 7 days in glycosylated buffered paraformaldehyde solution 30%. Afterward, the brains were frozen in isopentane at -80°C for posterior histological processing. All procedures were based on previous studies.

### Histology

Frozen brains were sliced in 50-μm coronal sections using a cryostat (Microm 550). Slices were arranged on gelatinized slides and then Nissl stained. A microscope (Zeiss Imager M2 ApoTome 2) under an objective lens of 20X was used to photograph individual brain sections on a mosaic through the Stereo Investigator R software (MBF Bioscience).

### Data analysis

All analyses were made using built-in and custom-made routines in MATLAB (MathWorks). We analyzed electrophysiological signals from (1) the running periods in which the treadmill belt was either under acceleration or constant speed, (2) the rest (intertrial interval) periods in which the treadmill was off (i.e., 0 cm/s). Signals from the first two seconds of the running periods were discarded to avoid potential cable torsion artifacts during the running initiation (i.e., due to the rat turning toward the running direction) and imprecise measures of the instantaneous treadmill speed during fast accelerations.

LFPs were filtered between 0.1 and 400 Hz and sampled at 1 kHz. One LFP channel from each session was selected based on the maximum ripple (150-250 Hz) band power. LFP signals were then visually inspected to exclude running trials with eventual saturation or mechanical artifacts. Time-frequency decompositions were made using the “spectrogram” function from the Signal Processing Toolbox (2-s window length with 90% overlap). Power spectral estimation was obtained using the “pwelch” function from the Signal Processing Toolbox (2-s window length with 25% overlap). To obtain band-pass filtered LFP signals, we used the “eegfilt” function (EEGLAB Toolbox, Delorme and Makeig, 2004). The instantaneous amplitude, phase, and frequency of a filtered signal were obtained using the “hilbert” function from the Signal Processing Toolbox.

We used the modulation index (MI) to measure the cross-frequency coupling between the phases of slow rhythms (0.5-12 Hz) and the amplitude of higher-frequency oscillations (20-100 Hz) in the LFPs (Tort et al., 2010). Briefly, the instantaneous phase of slow rhythms was split into 18 bins of 20° in which the average amplitude envelope of higher-frequency oscillations was measured. The MI values between multiple frequency pairs of slow and fast oscillations were computed and represented in color-code through the comodulogram, in which hot colors mean that the phase of the corresponding frequency in the x-axis modulates the amplitude of the frequency represented in the y-axis (for further details, see Tort et al., 2010 and Scheffer-Teixeira et al., 2012).

The phase-locking of spiking activity to delta and theta oscillations was measured by the mean resultant vector length (MVL) of the spike phases. The MVL for each unit was calculated from the average vector of all spike phases extracted from theta and delta filtered signals on a trial-by-trial basis. After this quantification, the MVL of each cell was considered the average for the trial type (1, 1.5, and 2 cm/s² and 20, 30, and 40 cm/s).

A naïve Bayes classifier was used to predict the running speed of the animal based on the LFP power spectra. We used a leave-one-out approach in which 1 to 12 Hz power spectra values (1-Hz step) across running speeds and trials (training bins) but one (test bin) were used to estimate the posterior probabilities. The classifier then predicts the running speed of the left-out power value at a given frequency. The percentage of correct decoding is obtained from the proportion of entries in the y=x diagonal of the confusion matrix of actual and decoded running speeds. The decoding performance was then compared to the distribution of 100 shuffled labels of running speed.

### Statistical analysis

Group results were expressed as mean ± SEM over trials or sessions, as indicated. We used one-sample t-tests, paired t-tests, and repeated-measures ANOVA followed by the Tukey-Kramer *post hoc* test to determine the statistical difference of group means, as appropriate. The Rayleigh test was used to compare phase distributions of spiking activity to a circular uniform distribution. Pearson correlation was used to estimate the linear relationship between delta and theta band power and peak frequency, as well as between the MVL for delta and theta phases. An alpha level of 0.05 was used to denote statistical significance. In the figures, one and two asterisks denote p<0.05 and p<0.01, respectively. All statistical analyses were performed in MATLAB.

## RESULTS

### Hippocampal delta oscillations emerge during stationary running

To investigate whether running speed modulates hippocampal delta oscillations, we trained 6 male adult Wistar rats to perform 48 treadmill runs at acceleration rates of 1, 1.5, and 2 cm/s² and runs at constant speeds of 20, 30, and 40 cm/s for 20 seconds with 5-s intertrial intervals of rest (Figure 1A and B). We used microdrives with 8 or 16 movable tetrodes to record local field potentials (LFPs) and spiking activity from the pyramidal layer of the dorsal CA1 area of the hippocampus (Figure 1C). Figure 1D shows a representative example of the instantaneous treadmill speed, raw and delta-filtered (0.5-3.5 Hz) LFPs, and associated spectral decomposition during a 5-s intertrial interval followed by a 20-s running at an accelerated speed protocol (2 cm/s²). Notice the emergence of delta oscillations with increasing running speed. Interestingly, for the constant speed protocols, normalized power spectra computed on a trial-by-trial basis revealed prominent shifts in the delta peak frequency across runs (Figure 1E and 1F). Additional examples of spectral decompositions of LFPs during runs at accelerated and constant speeds are shown in Figure 1-1.

We compared delta oscillations across runs at different constant speeds and across blocks of progressively increasing speeds. The top panel of Figures 2A shows the treadmill speed during runs at 20, 30, and 40 cm/s, while Figure 2B top panel shows the respective raw and delta band-filtered LFP signals. Average power spectra revealed a prominent increase in the delta band power and delta peak frequency with increases in running speed (Figure 2C top). Both delta band power and peak frequency were significantly higher during runs at 40 cm/s (Figure 2D and 2E bottom; p<0.05 and p<0.01 for band power and peak frequency, respectively, repeated-measures ANOVA followed by Tukey’s post hoc test, n=176 trials). We then compared delta oscillations across time blocks within runs at an accelerated speed protocol (2 cm/s²). The bottom panel of Figure 2A shows the treadmill speed divided into 3-s blocks (with associated speeds of 4-10, 22-28, and 34-40 cm/s), and Figure 2B bottom panel the respective raw and delta band-filtered LFP signals. Power spectra analysis showed a significant increase in the delta band power and delta peak frequency with faster speeds (Figure 2C, 2D, and 2E bottom panels; p<0.01 and p<0.05 for band power and peak frequency, respectively, repeated-measures ANOVA, n=176 trials).

**Figure 2.**
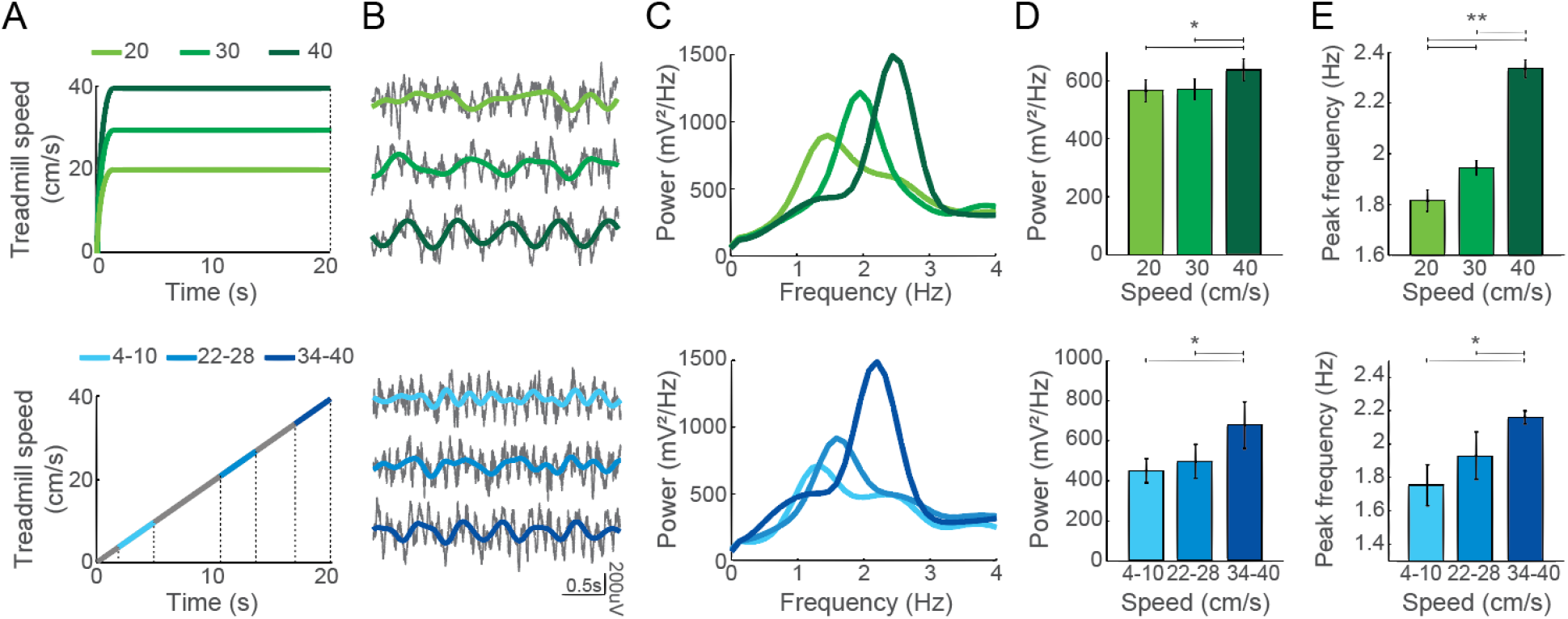
Higher running speeds are associated with higher delta power and peak frequency. (A) Schematic representation of the treadmill speed during running trials at 20, 30, and 40 cm/s (shades of green, top), and during a running trial at 2 cm/s² of acceleration (gray, bottom). For the latter, shades of blue indicate three 3-s blocks with speeds between 4-10 cm/s, 22-28 cm/s, and 34-40 cm/s, respectively. (B) Representative examples of raw (gray) and delta-filtered LFP signals (colored traces) obtained from three 3-s time blocks within the same trials as in A. (C) Group averaged power spectra in the delta band (same color code as in A). (D,E) Mean delta band power (D) and frequency (E) for constant (top) and accelerated (bottom) speeds. Bars denote mean and error bars ± SEM (*p<0.05 and **p<0.01, repeated-measures ANOVA followed by Tukey’s post hoc test, n=176 trials).

Since running speed has been often reported to correlate with hippocampal theta oscillations, we evaluated whether changes in delta band power and peak frequency could be explained by concomitant changes in the theta band. Spectrograms confirmed that delta and theta oscillations coexist in the hippocampus during treadmill running. Figure 3A shows representative running trials at 40 cm/s constant speed and 2 cm/s² of acceleration. Pearson’s correlation in 3-s blocks (n=3168 blocks) revealed no systematic relationship between the instantaneous delta and theta band power (Figure 3B; r = -0.05 for merged data from trials at 20, 30 and 40 cm/s of constant speed; r = 0.03 for merged trials at 1, 1.5 and 2 cm/s² of acceleration) nor between the instantaneous delta and theta peak frequency (Figure 3C; r = -0.06 for constant speed trials merged; r = 0.00 for accelerated speed trials merged).

**Figure 3.**
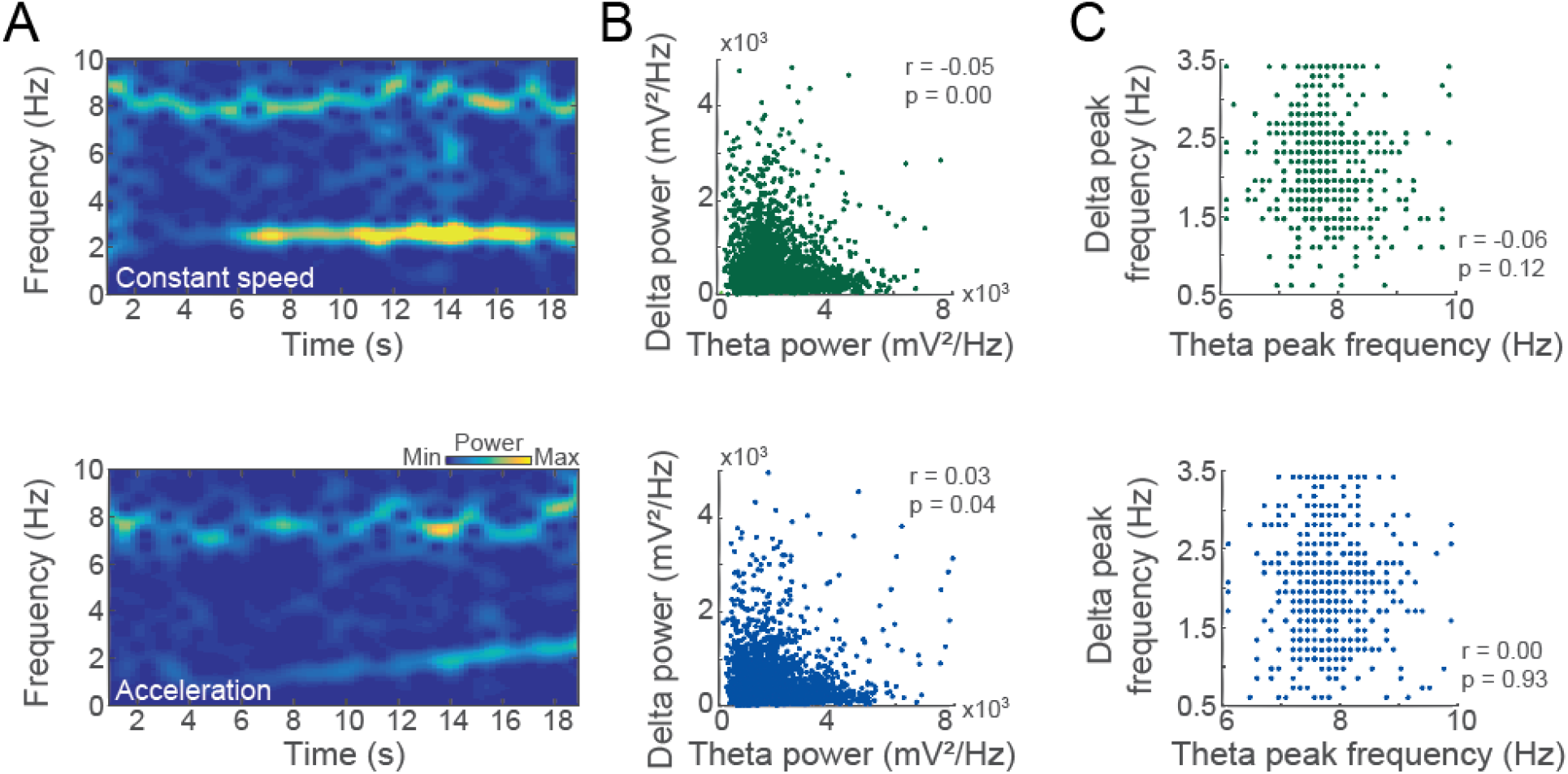
Changes in delta band power and peak frequency are not explained by changes in theta oscillations. (A) Spectrograms of representative running trials at a constant speed of 40 cm/s (top) and an accelerated speed (2 cm/s², bottom). (B) Scatter plot of the mean theta and delta band power across 3-s blocks for constant speed runs at 20, 30, and 40 cm/s (top) and accelerated runs at 1, 1.5, and 2 cm/s² (bottom, n=3168 blocks). (C) Scatter plot of the mean theta and delta peak frequency for the same time blocks as in B.

To investigate whether delta oscillations modulate the amplitude of higher frequency oscillations, we computed the phase-amplitude cross-frequency coupling (CFC) for the CA1 LFP. Figure 4A shows a representative example of CFC, in which the delta phase modulates the amplitude of low-gamma (LG, 20-50 Hz) but not the amplitude of the high-gamma (HG, 70-100 Hz) band. The maximal LG amplitude was locked to the peak of the delta wave (Figure 4A left and Figure 4B). We then compared CFC from signals recorded during runs at different constant speeds. Figure 4C shows the delta coupling strength for LG and HG during rest (black) and runs at 20, 30, and 40 cm/s (shades of green). We found that the strength of delta-LG and delta-HG coupling significantly increased with running speed (Figure 4D; p<0.01 for both delta-LG and delta-HG coupling, repeated-measures ANOVA followed by Tukey’s post hoc test, n=22 sessions from 6 rats).

**Figure 4.**
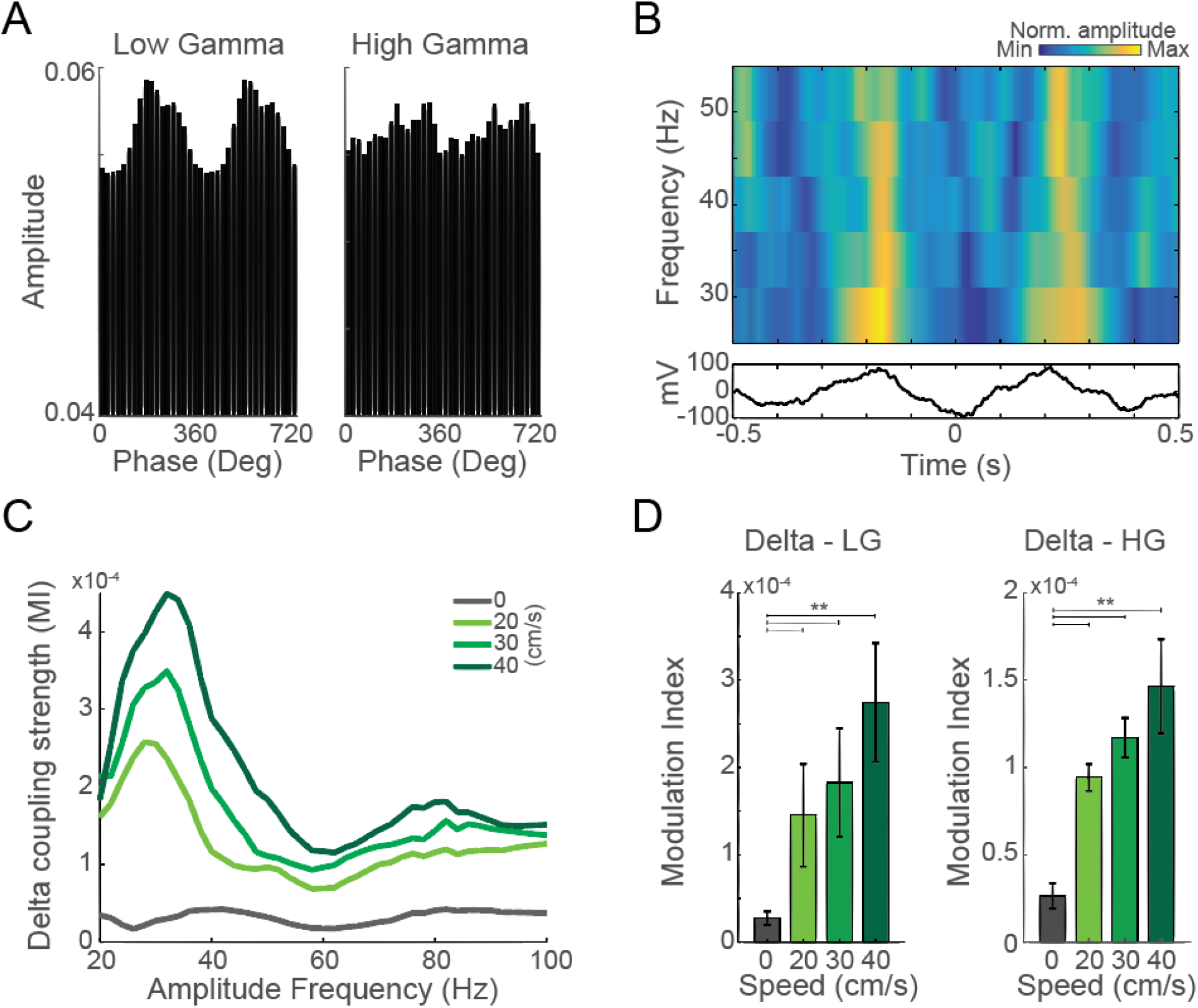
Delta phase modulates the amplitude of gamma oscillations. (A) Cross-frequency coupling between the phase of delta (1.5-3.5 Hz) and the amplitude of two gamma sub-bands computed using data from the whole session (48 runs and intertrial intervals; LG: 20-50 Hz and HG: 70-100 Hz). (B) Time-frequency amplitude distribution triggered by the delta trough shows maximal LG and HG amplitude at the peak of the delta wave. (C) Mean coupling strength between the phase of delta (1.5-3.5 Hz) and the amplitude of gamma (20-100 Hz) during runs at different speeds, as labeled. (D) Mean modulation index between delta phase and the amplitude of LG (left) or HG (right) during runs at constant speeds. Bars denote mean and error bars ± SEM (**p<0.01, repeated-measures ANOVA followed by Tukey’s post hoc test, n=22 sessions).

To determine whether delta oscillations entrain the spiking activity of hippocampal neurons, we next measured the spike-phase coupling of putative pyramidal cells and interneurons during treadmill runs at constant speeds. Figure 5A shows the spiking activity of a representative pyramidal cell simultaneously locked to delta and theta phase (left) and a representative interneuron locked to theta but not delta phase (right). At the group level, we found that 54.7% of the pyramidal cells significantly phase-locked to delta oscillations (52 from 95 units, p<0.05, Rayleigh’s test for circular uniformity), and that 26.3% (25 units) simultaneously phase-locked to delta and theta oscillations. In contrast, 10.2% of the interneurons were phase-locked to delta oscillations (5 from 49 units), and only 4.1% (2 units) were simultaneously phase-locked to delta and theta (Figure 5 and Figure 5-3).

**Figure 5.**
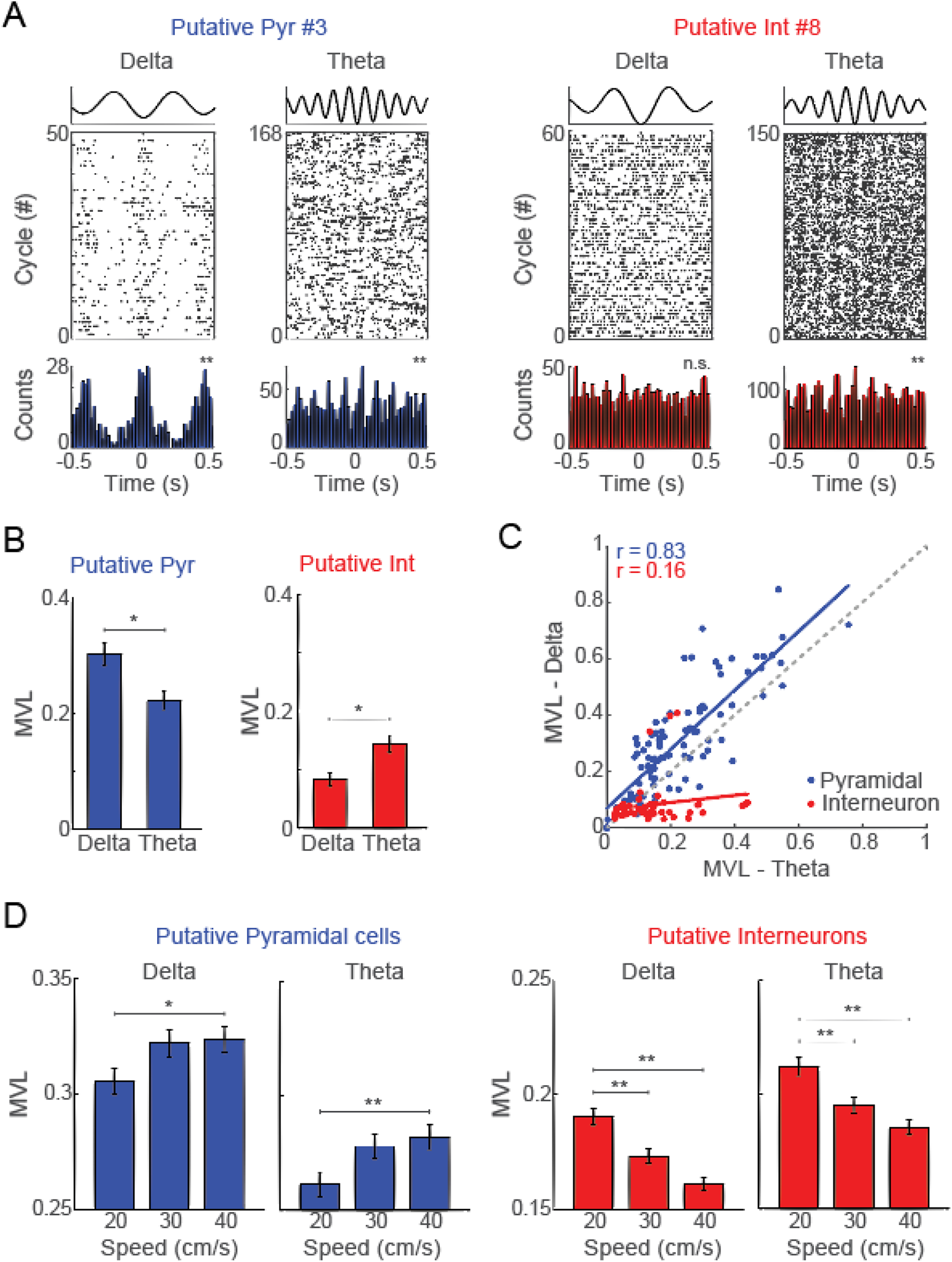
Delta oscillations entrain pyramidal cells but not interneuron spiking activity. (A) Representative examples of spiking activity from a putative pyramidal cell (left) and an interneuron (right) within delta and theta cycles. (B) Mean vector length (MVL) of pyramidal cells (left) and interneurons (right) for delta and theta phases. (C) Scatterplot of delta and theta MVL for pyramidal cells (blue dots) and interneurons (red dots). Gray dots represent the MVL of unclassified cells. Linear regressions for pyramidal cells and interneurons are fitted as blue and red lines, respectively. (D) Pyramidal cell (left) and interneuron MVL (right) for delta and theta phase at different running speeds. Bars denote mean and error bars ± SEM (*p<0.05 and **p<0.01, repeated-measures ANOVA followed by Tukey’s post hoc test).

Comparing the mean vector length (MVL) for delta vs. theta, we surprisingly found that pyramidal cells coupled stronger to delta phase, while most interneurons coupled stronger to theta phases (Figure 5B; p<0.05, paired t-test). The Pearson correlation coefficient between delta and theta MVL was r = 0.83 for pyramidal cells and r = 0.16 for interneurons (Figure 5C), where pyramidal cells are more likely to be coupled to both rhythms while interneurons are most coupled to theta oscillations. The conjunctive analysis of spike coupling to delta and theta phases revealed that pyramidal neurons were simultaneously entrained by both oscillations, while interneurons were mostly entrained by theta phases (Figure 5-4).

We next compared the MVL of pyramidal cells and interneurons during treadmill runs for different trials at constant speeds (20, 30, and 40 cm/s). We found that pyramidal neurons significantly increased their coupling strength with running speed while the spike-phase coupling strength of interneurons decreased (Figure 5D, p<0.05 and p<0.01, repeated-measures ANOVA followed by Tukey’s post hoc test).

Next, we used a Bayesian decoder to classify running speed based on spectral decompositions of the LFP (power values from 1 to 12 Hz). We first classified running speed during constant speed protocols (20, 30, and 40 cm/s) and found the decoder performance to peak at 2 Hz and 8 Hz (Figure 6A top). The decoding performance using oscillatory frequencies at 2 Hz was higher than 99% (3 SDs) of 100 surrogated data (p<0.01, one-sample t-test against 0.33); moreover, the performance at 2 Hz was significantly higher than at 8 Hz (Figure 6B top, p<0.01, paired t-test, n=22 sessions). For accelerated speed protocols, the decoding performance (six 3-s time blocks) also peaked at 2 Hz (Figure 6A bottom). At this frequency, decoding performance was higher than that of 95% (2 SDs) of 100 surrogated data, while at 8 Hz the decoding performance did not differ from chance (p<0.01 and p=0.81, respectively, one-sample t-test against 0.17, n=22 sessions). Moreover, decoding performance was significantly higher at 2 Hz than at 8 Hz (Figure 6B bottom, 2 vs. 8 Hz, p<0.01, paired t-test, n=22 sessions).

**Figure 6.**
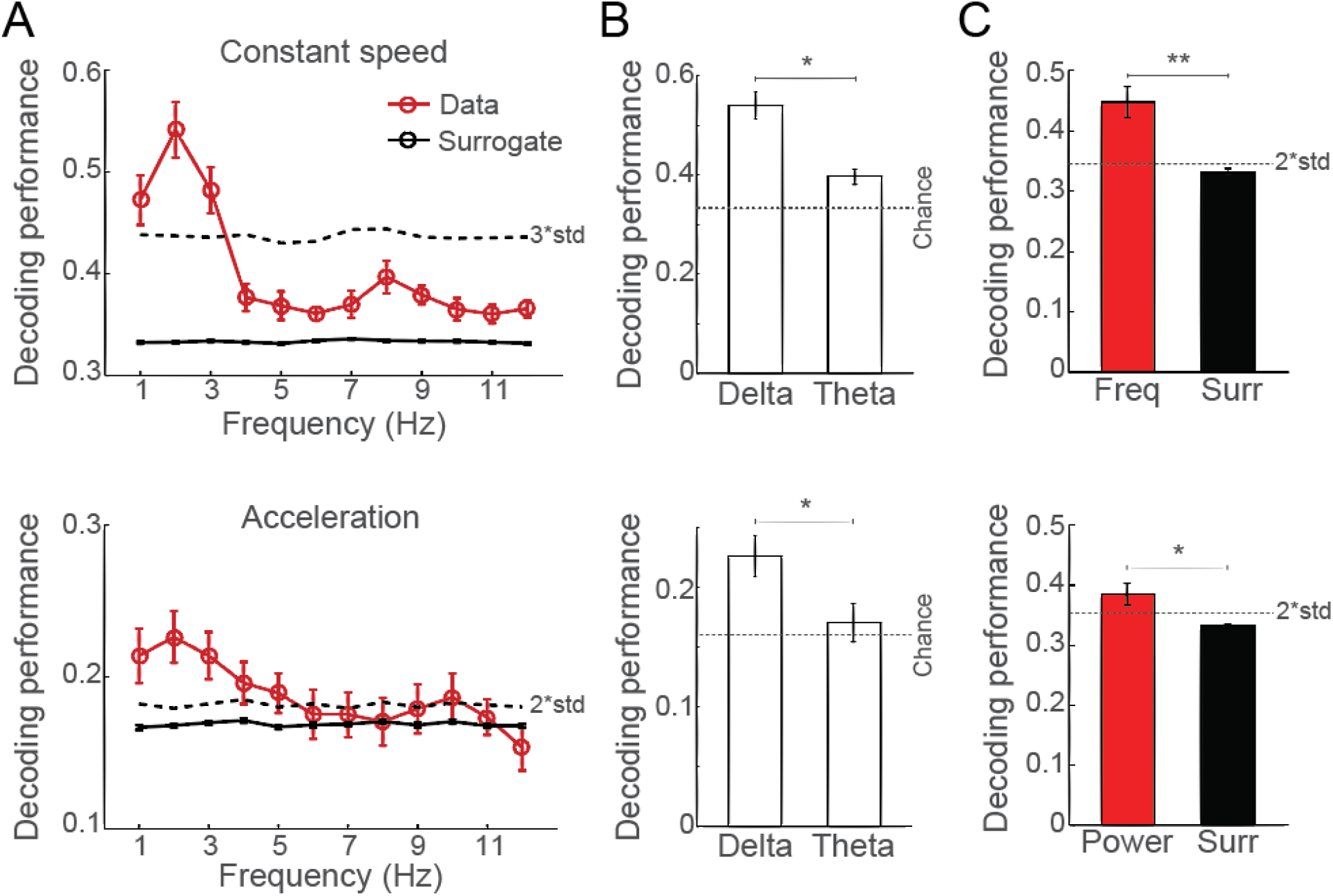
Delta oscillations predict running speeds. (A) Decoding performance for runs at constant (20, 30, and 40 cm/s, top) and accelerated speeds (2 cm/s², bottom). Mean classification based on the actual power spectra are shown in red and the mean surrogated data in black (n=22 sessions). Dashed lines show 3 and 2 times the SD of the surrogate distributions for runs at constant and accelerated speeds, respectively. (B) Average decoding performance for the delta and theta band during runs at constant (top) and accelerated (bottom) speeds (*p<0.05 for delta vs. theta, paired t-test). The dotted line shows the decoding probability expected by chance (0.33 and 0.16 for constant and accelerated speeds, respectively). (C) Average decoding performance using the delta peak frequency (top) and peak power (bottom) for runs at constant speeds. Bars denote mean and error bars ± SEM. Black dashed lines depict 2 times the SD of the surrogate distribution (*p<0.05 and **p<0.01, paired t-test).

Finally, we independently fed the classifier with peak frequency or peak power values at the delta band (Figure 6C top and bottom, respectively), and found that both parameters could predict speed better than chance (p<0.01 and p<0.05 for peak frequency and peak power, respectively, one-sample t-test against 0.33, n=22 sessions). The direct comparison showed that delta peak frequency had higher decoding performance than delta peak power (p<0.01, paired t-test, n=22 sessions).

## DISCUSSION

In the current study, we demonstrate the emergence of prominent oscillatory activity in the delta frequency (∼2 Hz) in the hippocampus of rats performing stationary runs on a treadmill. Running speed increased delta power and frequency in runs at constant and accelerated speeds (Figures 1 and 2). Spectral changes in the delta band were independent of changes in the theta band (Figure 3). Moreover, the delta phase modulated the amplitude of low-gamma in a speed-dependent way (Figure 4) and entrained spiking activity of putative pyramidal cells (Figure 5). Finally, delta power could predict running speed using a naïve Bayes classifier (Figure 6).

Running speed has been previously correlated with the amplitude and frequency of hippocampal theta and gamma rhythms (Ahmed and Mehta, 2012). Delta oscillations are instead usually reported as absent in the hippocampus of rats engaged in locomotor activity (Chrobak and Buzsáki, 1994; Schultheiss et al., 2020). Prominent delta power has been often observed during quiet behaviors and slow-wave sleep, which exhibit no theta rhythm (Emmons et al., 2017; Hirase et al., 2001; Khazipov and Luhmann, 2006; Siapas and Wilson, 1998; Sirota et al., 2003; Steriade et al., 2001; Sirota and Buzsáki, 2005). However, contrary to this classical dichotomy between theta versus delta hippocampal states, we show here the concomitant occurrence of both oscillations during stationary running on a treadmill. Importantly, neither their instantaneous power nor their instantaneous frequency was correlated, indicating the simultaneous expression of two distinct and independent oscillatory activities within the 1-10 Hz frequency range. Since many studies apply the theta/delta power ratio to classify epochs of immobility and exploratory states (Csicsvari et al., 1999; O’Neill et al., 2006; Korotkova et al., 2010; Penley et al., 2011; Schultheiss et al., 2020), our results suggest caution and a more careful inspection of the delta band activity, at least when analyzing experiments involving stationary running.

There is, however, provocative new evidence in the literature that associates hippocampal delta oscillations and stationary locomotion. Namely, we recently reported a progressive increase of delta power across consecutive runs at the same speed on a treadmill, which points to rhythmic delta activity as a potential biomarker of running-associated fatigue (Furtunato et al., 2020). Here, we advance this idea by showing that both delta power and frequency increase with increases in stationary running speed on a treadmill (Figures 1 and 2). In line with our findings, preliminary indicative data of speed-modulated delta power have also been shown in running wheels, though these results were not further explored (Czurkó et al., 1999; Molter et al., 2012). More recently, the amplitude of 4-Hz oscillations was also associated with running speed in a virtual reality apparatus that resembles stationary locomotion (Safaryan and Mehta, 2021). The underlying mechanisms responsible for generating locomotion-related delta oscillations specifically in stationary running conditions remain unknown.

To further evaluate how delta oscillations relate to other hippocampal rhythms, we analyzed whether the delta phase modulates the amplitude of faster rhythms (Buzsáki and Draguhn, 2004; Jensen and Colgin, 2007; Penttonen and Buzsáki, 2003; Tort et al., 2010, 2008). Despite the concomitant occurrence of delta and theta oscillations, we found no evidence of phase-amplitude cross-frequency coupling between these bands (not shown). Instead, we found that the delta phase modulates the amplitude of higher-frequency oscillations circumscribed into the low-gamma sub-band (20-55 Hz). The maximal low-gamma activity coupled to the peak of the delta wave (Figure 4), similarly to the theta-gamma phase-amplitude coupling (Scheffer-Teixeira et al., 2012; Schomburg et al., 2014). It should be noted, however, that the delta-low gamma coupling strength was lower (MI∼10^-4^) than traditionally observed for theta-gamma interactions (MI ∼10^-2^ - 10^-3^; Scheffer-Teixeira and Tort, 2018). These resemblances and differences suggest that delta oscillations are not segregated from other oscillatory components but interact with coexisting rhythms in the rat hippocampus. Moreover, the speed-dependence of the delta-LG coupling strength highlights the role of the delta phase in the organization of other hippocampal rhythms during treadmill running. Of note, while previous studies reported that theta phase modulates the amplitude of both low- and high-gamma oscillations in a speed-dependent way (Chen et al., 2011; Sheremet et al., 2019), to the best of our knowledge, no previous report has shown delta-gamma phase-amplitude coupling in the rodent hippocampus during active locomotor behaviors.

The firing rates of CA1 neurons increase with running speed during locomotion through the environment and during stationary running (Czurkó et al., 1999; Montgomery et al., 2009; Ahmed and Mehta, 2012; Góis and Tort, 2018; Iwase et al., 2020). Here, we observed that putative interneurons increased firing rates with running speed (Figure 5-1; Figure 5-2) and that their activity was predominantly coupled to theta oscillations and only marginally affected by the delta phase (Figures 5 and 5-3). In contrast, 54.7% of the pyramidal cells were phase-locked to delta oscillations; moreover, pyramidal cells exhibited even higher coupling strength to delta than to theta phase (Figure 5). Additionally, the coupling strength of pyramidal neurons to delta phase increased at faster running speeds, suggesting that the emergence of delta oscillations entrained the rhythmicity of pyramidal neurons in a speed-dependent way. These results differ from previous findings from Safaryan and Mehta (2021) that observed high levels of coupling between interneurons and 4-Hz oscillations, while pyramidal neurons were only weakly entrained when rats ran in a virtual reality apparatus. Nevertheless, taken together, these data show that oscillations at the delta frequency coordinate the rhythmicity of spiking activity of hippocampal neurons during certain types of locomotor behavior.

Finally, we asked whether locomotion-related delta oscillations carry information about running speed. By using a naive Bayes classifier, we could successfully decode running speed from power spectra values from 1 to 12 Hz. Surprisingly, the decoding performance was higher using the oscillatory activity at the delta (∼2 Hz) than theta (∼8 Hz) band, which is remarkable since theta oscillations are usually reported as a strong predictor of running speed during translational and stationary conditions (McFarland et al, 1975; Sławińska et al, 1998; Richard et al, 2013; Li et al, 2014; Bender et al, 2015; Safaryan and Mehta, 2020). Moreover, the fact that delta frequency was better than delta power to classify running speed highlights that different oscillatory parameters in the delta band may convey different amounts of information about running speed (Figure 6). Whether delta power and frequency could be independently entrained by particular sets of physiological conditions remains to be investigated. As far as we know, this is the first study to show that oscillatory activity at the delta band can robustly predict running speed.

In all, we conclude that delta oscillations, in coordination with other rhythms, may underlie the processing of locomotion speed in the hippocampus of rats performing stationary running in the presence of static external landmarks.

## Acknowledgments

This work was supported by the Conselho Nacional de Desenvolvimento Cientifico e Tecnológico (CNPq) and Coordenação de Aperfeiçoamento de Pessoal de Nível Superior (CAPES). We would like to thank Richardson Leão, George Nascimento, and Izabela Paiva for technical assistance.

## Author contributions

A.M.B.F., R.P., and H.B. designed research; A.M.B.F. and R.P. performed experiments; A.M.B.F., R.P., and H.B. analyzed data; A.M.B.F., R.P., B.L.S., A.B.L.T, and H.B. wrote the manuscript.

## EXTENDED DATA

**Figure 1-1.**
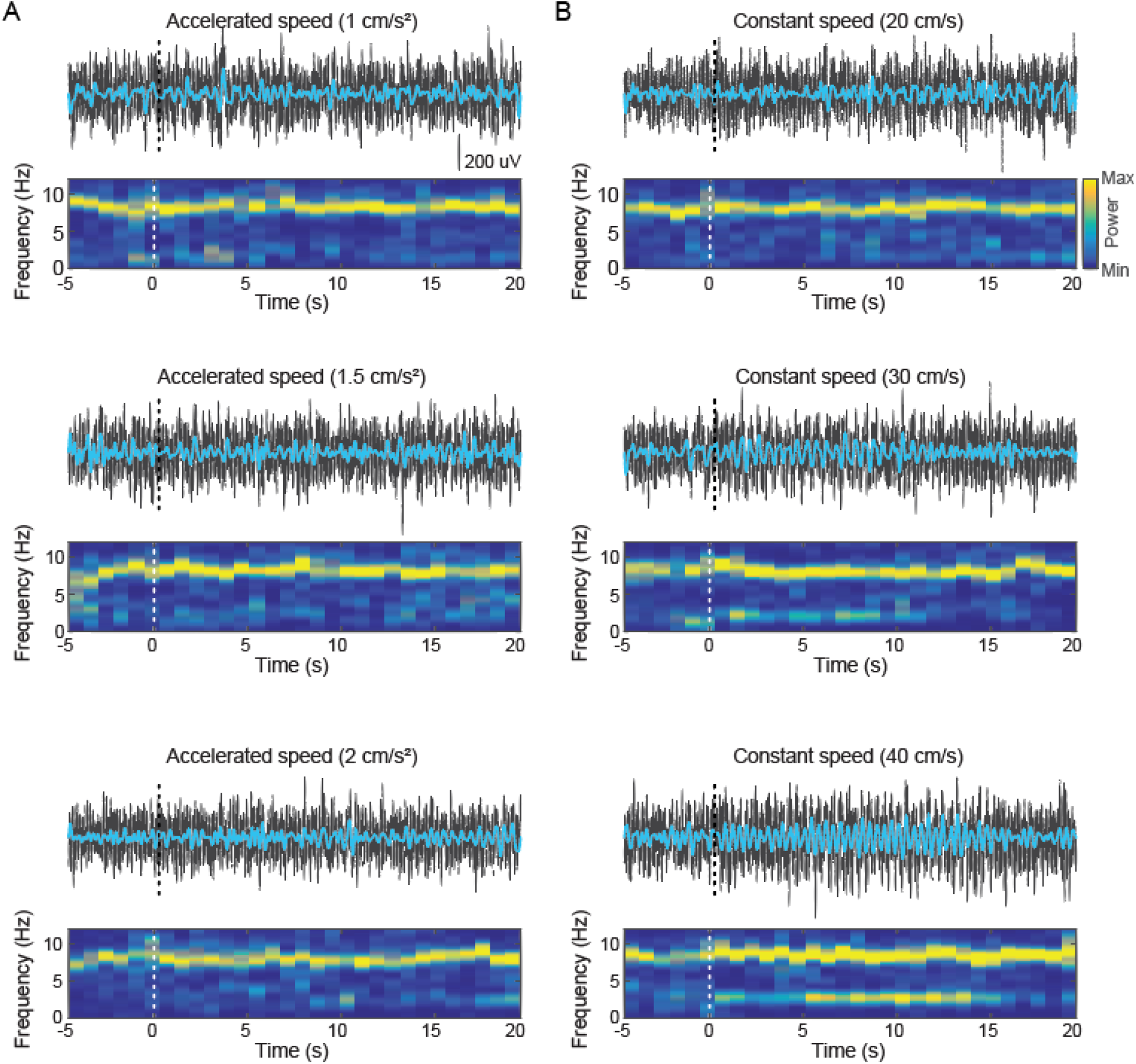
Raw (black) and delta-filtered (cyan) LFP signals and the associated spectrograms (0-12 Hz) during representative runs at accelerated (A) and constant speeds (B). Each panel shows the 5-s intertrial interval and the following 20-s treadmill running. Dashed lines depict the time the treadmill started. In addition to the prominent theta activity, notice oscillatory activity in the delta frequency band (∼2 Hz) at the higher running speeds.

**Figure 5-1.**
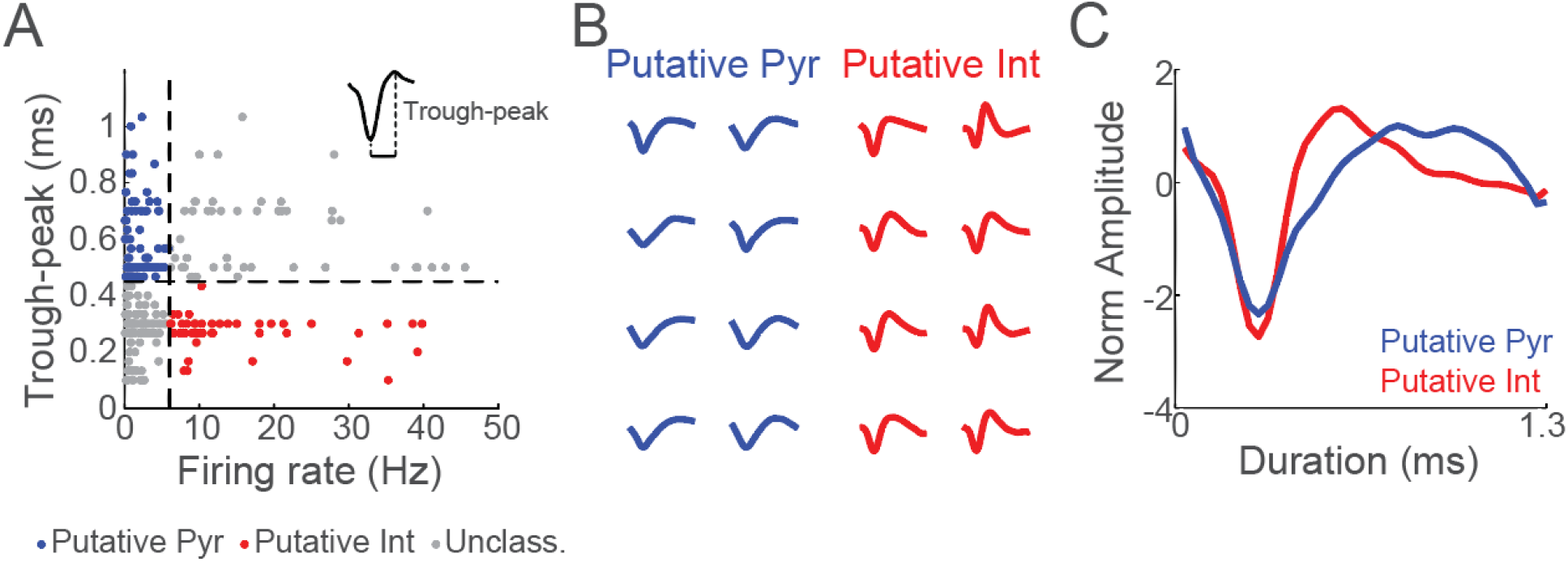
Neuron type classification. (A) Scatter plot of the trough to peak duration and firing rate of neurons. Blue, red, and gray dots represent putative pyramidal neurons, interneurons, and unclassified neurons. The inset shows the duration of the trough-to-peak interval. (B) Representative examples of spike waveforms classified as putative pyramidal units (blue) and putative interneurons (red). (C) Normalized spike amplitude of putative pyramidal and interneuron units.

**Figure 5-2.**
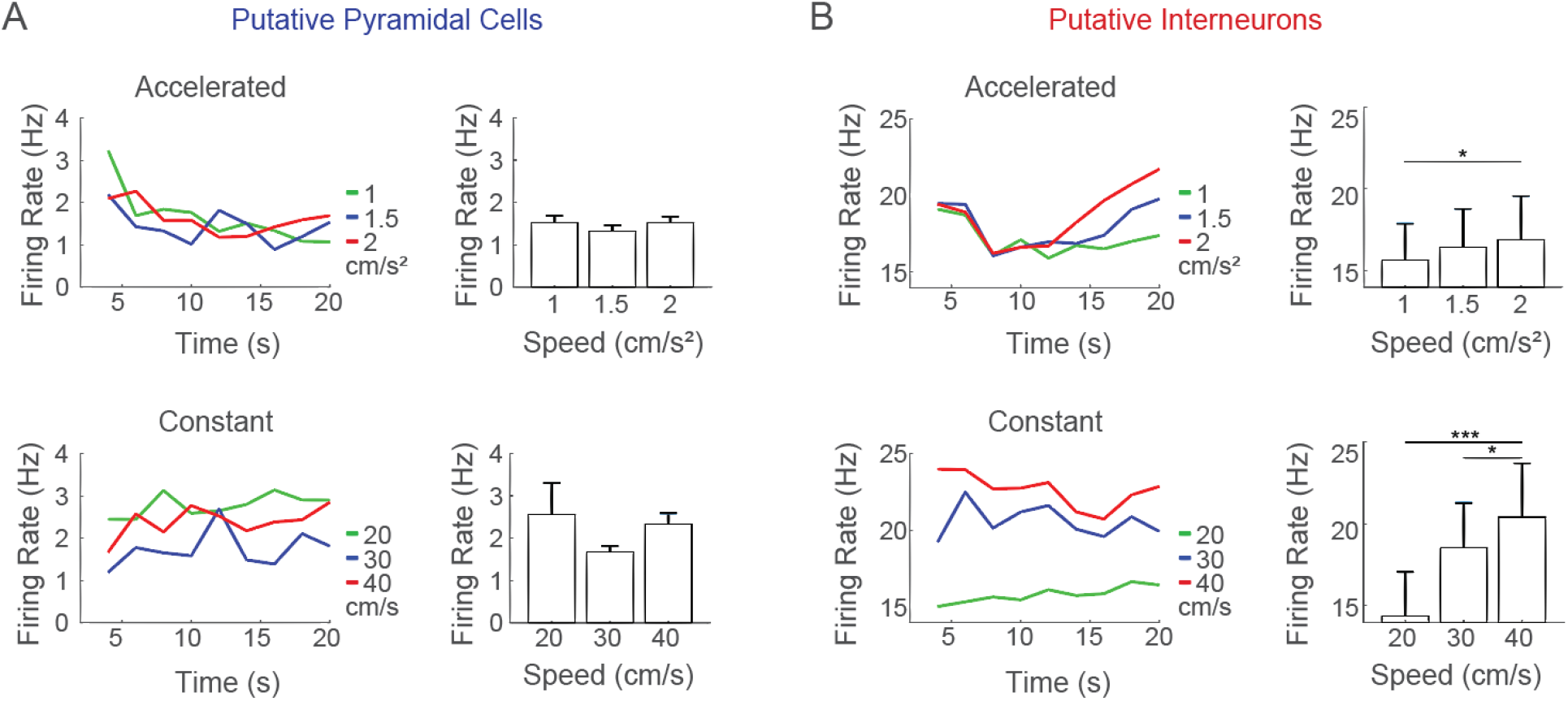
Mean firing rates of putative pyramidal cells (A) and interneurons (B) during treadmill runs at accelerated (upper) and constant speeds (lower).

**Figure 5-3.**
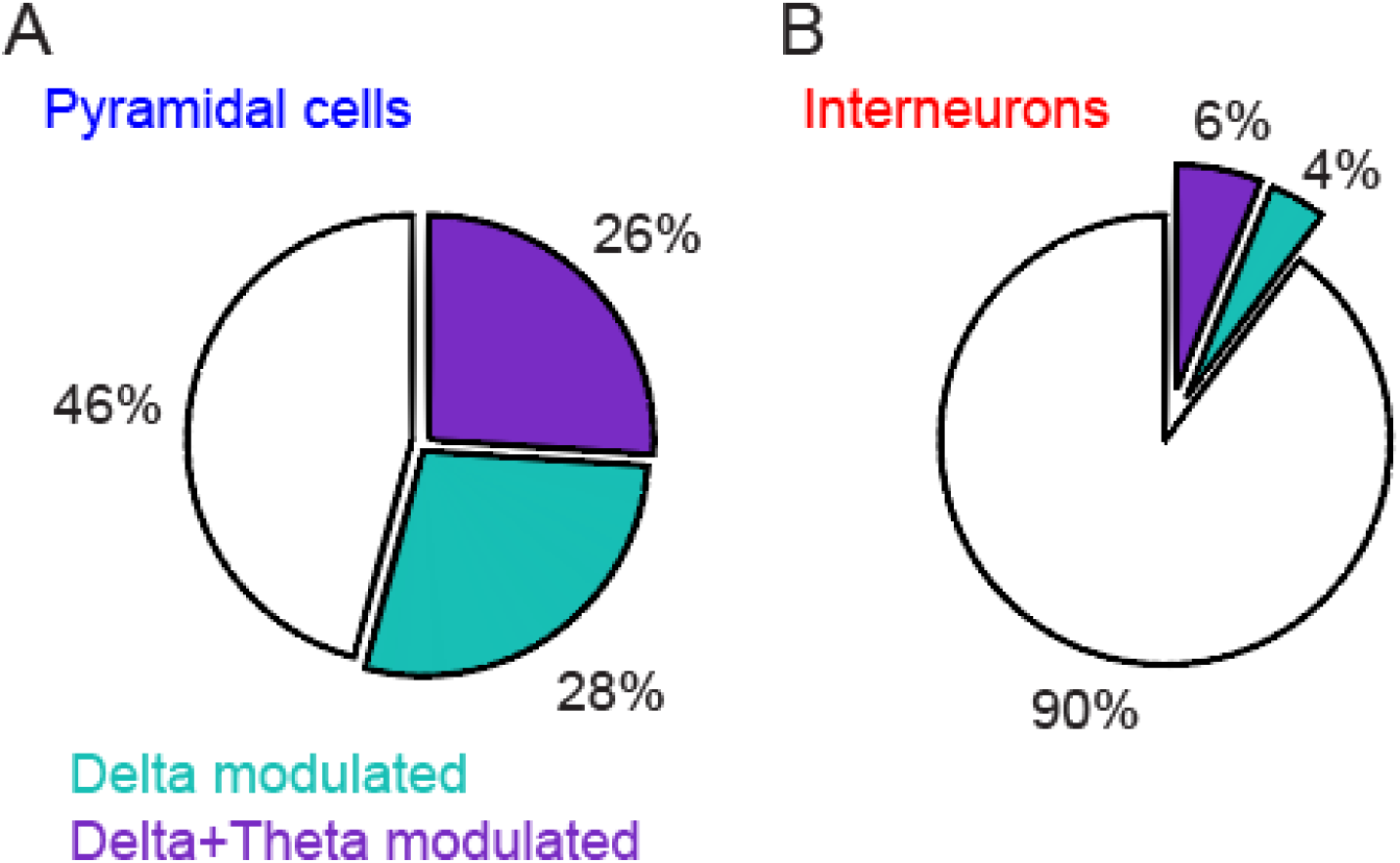
The proportion of putative pyramidal cells and interneurons locked to delta and theta phases. (A) 54% of the pyramidal cells were coupled to the delta phase (green + purple), and 26% simultaneously locked to the delta and theta phases (purple). (B) 10% of the interneurons were coupled to the delta phase (green + purple), and 6% simultaneously coupled to the delta and theta phases (purple).

**Figure 5-4.**
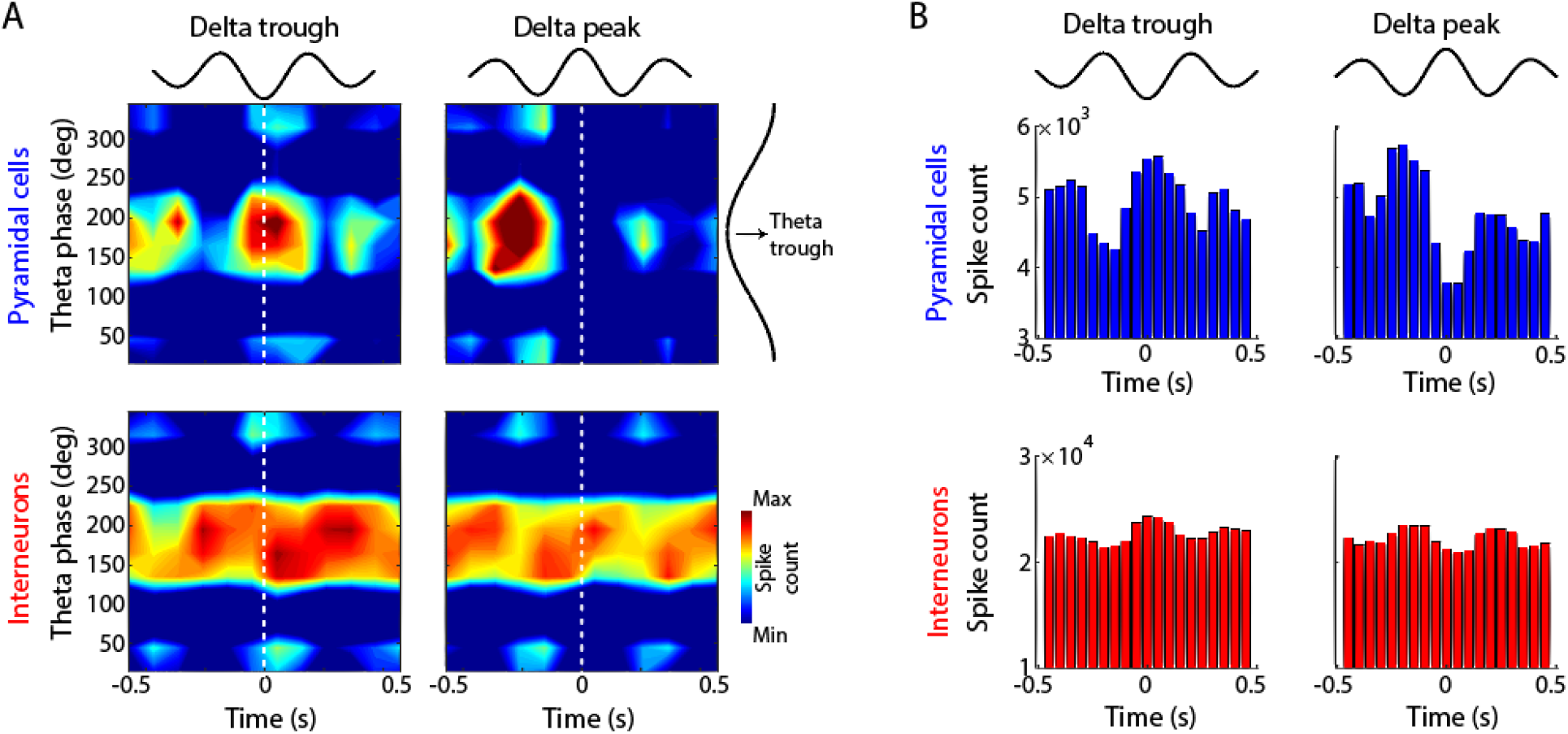
Joint modulation of putative pyramidal cells and interneurons by delta and theta phases. (A) Spike distribution map aligned to delta and theta phases during treadmill runs at constant (40 cm/s) speed. Notice that most spikes of pyramidal cells were simultaneously locked to delta and theta troughs (top), while spikes of interneurons were modulated by theta phases (bottom). (B) Pyramidal cells’ and interneurons’ spike modulation around delta trough and peak.

